# Evaluating a Novel High-Density EEG Sensor Net Structure for Improving Inclusivity in Infants with Curly or Tightly Coiled Hair

**DOI:** 10.1101/2024.03.18.584988

**Authors:** Nwabisa Mlandu, Sarah A. McCormick, Lauren Davel, Michal R. Zieff, Layla Bradford, Donna Herr, Chloë A. Jacobs, Anele Khumalo, Candice Knipe, Zamazimba Madi, Thandeka Mazubane, Bokang Methola, Tembeka Mhlakwaphalwa, Marlie Miles, Zayaan Goolam Nabi, Rabelani Negota, Khanyisa Nkubungu, Tracy Pan, Reese Samuels, Sadeeka Williams, Simone R. Williams, Trey Avery, Gaynor Foster, Kirsten A. Donald, Laurel J. Gabard-Durnam

## Abstract

Electroencephalography (EEG) is an important tool in the field of developmental cognitive neuroscience for indexing neural activity. However, racial biases persist in EEG research that limit the utility of this tool. One bias comes from the structure of EEG nets/caps that do not facilitate equitable data collection across hair textures and types. Recent efforts have improved EEG net/cap design, but these solutions can be time-intensive, reduce sensor density, and are more difficult to implement in younger populations. The present study focused on testing EEG sensor net designs over infancy. Specifically, we compared EEG data quality and retention between two high-density saline-based EEG sensor net designs from the same company (Magstim EGI, Whitland, UK) within the same infants during a baseline EEG paradigm. We found that within infants, the tall sensor nets resulted in lower impedances during collection, including lower impedances in the key online reference electrode for those with greater hair heights and resulted in a greater number of usable EEG channels and data segments retained during pre-processing. These results suggest that along with other best practices, the modified tall sensor net design is useful for improving data quality and retention in infant participants with curly or tightly-coiled hair.

---

Electroencephalography, or EEG, is an integral tool for recording brain activity noninvasively in humans. This electrophysiological approach can be used in both clinical and research settings and provides researchers and clinicians with valuable information with regards to brain functioning. For example, in clinical practice, it is commonly used to diagnose and monitor several brain disorders, such as epilepsy (Benbadis et al., 2020). In research, EEG can be used to measure specific patterns of electrical activity associated with sensory, motor, affective, and cognitive events, to track brain development and maturation, and to describe brain-behaviour relations (e.g. Beres, 2017; Carbine et al., 2018; Clayson et al., 2021; Jones et al., 2016; Remijn et al., 2014, Kaiser et al., 2020).

Although decades of research and clinical practice support the potential of EEG for understanding and treating the human condition, this tool’s utility has been significantly limited because EEG research has historically systematically limited inclusion of participants with curly or tightly coiled hair, particularly those of African or African American descent. Numerous studies have described such systemic racial biases in EEG research (e.g., Andrews & Swaine, 2022; Garcini et al., 2022; Girolamo et al., 2022; Green et al., 2022; Norton et al., 2021; Penner et al., 2022; Webb et al., 2022). One core reason accounting for this exclusion is that the EEG sensors and cap/net structures are not designed with these populations’ hair textures in mind. In order to record valid and reliable data for the analysis of EEG, it is essential for the electrodes to properly adhere to the scalp and reduce extraneous noise and resistance (Girolamo et al., 2022). Current cap and net structures meant to fit close to the scalp with short-profile electrodes make obtaining an adequate, reliable signal challenging for those with curly and tightly coiled hair. Curly or tightly coiled hair can be left natural or is often developed into various hairstyles such as braids, cornrows or locs. With current cap/net structures, these hair types and styles have a high probability of impeding adherence to the scalp and thus impeding recording quality EEG data.

This exclusionary EEG equipment design has ramifications from collection to analysis of EEG data. Specifically, some EEG labs may consider hairstyles like braids, cornrows, and locs, or curly or tightly coiled hair textures to be exclusionary for participation in studies due to anticipated poor adherence between the scalp and the electrodes (Choy et al., 2022). Therefore, even before the data collection has ensued, there may be guidelines instituted that implicitly or overtly exclude mainly people of colour of African descent. In the case where EEG research includes people of color and African descent with curly or tightly coiled hair, data retention remains an issue. For instance, in some cases where a saline solution or gel has been used for better conductivity, the electrode impedances may still remain high due to poor absorption, producing a poor signal (Norton et al., 2021). Moreover, the EEG cap itself may not be a good or tight fit due to the hair being thick and/or tall, thus the impedances between electrodes and skin may remain high. The acceptance and usability of the collected data for statistical analysis is commonly dependent on strict impedance thresholds and signal-to-noise ratios (Parker & Ricard, 2022). As a result, the data collected from these participants may not be usable due to significant extraneous noise and higher impedances. Consequently, fewer participants of color or African descent with tightly coiled hair textures may be represented in the datasets of these research studies. These exclusionary practices greatly reduce the generalizability of EEG findings due to a lack of diversity in these studies and impede asking some critical research questions altogether (Norton et al., 2021).

A few studies have presented solutions for this problem in adults. For instance, Etienne et al. (2020) proposed the use of a newly developed innovative electrode, *Sevo*, and braiding the hair into cornrows to reduce impedances and improve the quality of EEG data from people with coarse and curly hair. This solution is advantageous in many respects but as with most adult EEG systems, it requires significant time to correctly apply the electrodes, and it may be best suited for lower-density EEG montages (often 32 or fewer electrodes). Given more limited participant tolerance and attention and wake spans in early development, this novel method may not generalize across the lifespan. However, there have been a dearth of studies that have described the problem or proposed solutions to this issue over infancy or early childhood. Therefore, we aim to first establish the scope of the problem with respect to participant exclusion, particularly for infants with coarse, curly or tightly coiled hair, by evaluating a widely-used EEG system in infancy, the Magstim EGI 128-channel HydroCel Geodesic Sensor Net and saline based system. Second, we have partnered with Magstim EGI to test and evaluate a new sensor net design with taller electrodes as a potential solution. We conduct these assessments within the context of Khula, a longitudinal study on infant neurodevelopment taking place in Cape Town, South Africa. Finally, we make recommendations for future infant research and EEG equipment design to improve inclusion of participants with all hair types in electrophysiological research.

## 2. Method

### 2.1 Participants

The initial sample consisted of 6- to 11-month-old infants (*N* = 47) drawn from the Khula study, a larger longitudinal study on infant development (Zieff et al., 2024). Families were recruited for the larger study from an established pregnancy registry site at the Gugulethu Midwife Obstetrics Unit, located in Gugulethu (an informal urban settlement in the Cape Town metropole in South Africa). Women who were pregnant, in their third trimester of pregnancy (28 - 36 weeks) and over the age of 18 years at the time of recruitment were eligible to participate in the larger study. Exclusion criteria from the larger study included multiple pregnancy, psychotropic drug use during pregnancy, congenital malformations and abnormalities (e.g., spina bifida, Down’s syndrome), and significant delivery complications (e.g., uterine rupture, birth asphyxia). Participants were excluded from analyses in the current study based on inconsistent net placement or excessive fussiness during EEG recordings (*N* = 12 infants). The final sample in this study included 35 infants (*N* = 14 females; *M*_*age*_ = 252.97 days, *SD* = 37.12 days). Participant sociodemographics are reported (Table 1). Families were compensated for their time, in line with local Human Research Ethics guidelines and free transportation was provided to all families. All procedures were approved by the University of Cape Town Institutional Review Board.

**Table 1.** Demographic information reported by caregivers and collected during study sessions. ZAR = South African Rand (at the time of writing, USD1.00 = ZAR19.00).

|  | No. (%) with data |
| --- | --- |
| **Maternal education** | 35 (100%) |
| Completed Grade 8 to Grade 11 | 18 (51%) |
| Completed Grade 12 | 12 (34%) |
| Part of university/college/post-matric education | 4 (11%) |
| Completed university/college/post-matric education | 1 (3%) |
| **Language spoken in home (primary)** | 35 (100%) |
| Xhosa | 34 (97%) |
| Zulu | 1 (3%) |
| **Language spoken in home (secondary)** | 14 (40%) |
| English | 12 (34%) |
| Another language (e.g. Sotho, Afrikaans, Shona) | 2 (6%) |
| **Household income** | 35 (100%) |
| Less than ZAR1000 per month | 6 (17%) |
| ZAR1000-ZAR5000 per month | 21 (60%) |
| ZAR5000-ZAR10,000 per month | 7 (20%) |
| Unknown | 1 (3%) |
| **Hair height (cm)** | 35 (100%) |
| 0 cm ≤ 0.5 cm | 7 (20%) |
| > 0.5 cm ≤ 1 cm | 7 (20%) |
| > 1cm | 21 (60%) |

### 2.2. Hair height measurement

We measured the length, or height, of the infants’ hair as it lay naturally on the scalp. Hair height was measured in centimeters (cm), and was measured prior to the placement of any net or the application of conditioner. Hair height was generally measured around the vertex of the scalp, or where the hair appeared to be the tallest and/or most dense. Hair height was logged for all infants using the following categories: 1) 0 cm ≤ 0.5 cm, 2) > 0.5 cm ≤ 1 cm, 3) > 1 cm.

### 2.3 Procedure and EEG Data Acquisition

The goal of the current study was to compare the effectiveness of two different types of net structures, both manufactured by Magstim Electrical Geodesic Inc (Magstim EGI; Whitland, UK), in collecting high-quality EEG data with infants who have curly or tightly coiled hair. Both nets were versions of the 44-47 cm infant sized 128-channel HydroCel Geodesic Net (Magstim EGI). All infants included in testing had head circumference measurements that fell within the range for use with this net (*M* = 45.77 cm, *SD* = 1.18 cm, range: 43.5 - 47.5 cm). The short sensor net is the standard net structure currently available for purchase through Magstim EGI. In this standard short net, all of the pedestals in the net are the same 6.5 mm height encased with a soft plastic material, and the tall sensor net includes a large section of electrode pedestals that are 9.3 mm tall where hair is likely to be present with a rigid plastic encasing to maintain the structure (Figures 1 - 3). The tall net structure was designed to hopefully better accommodate different structures and volumes of hair, but still be easy to place in infancy when there is less time to place individual electrodes.

**Figure 1:**
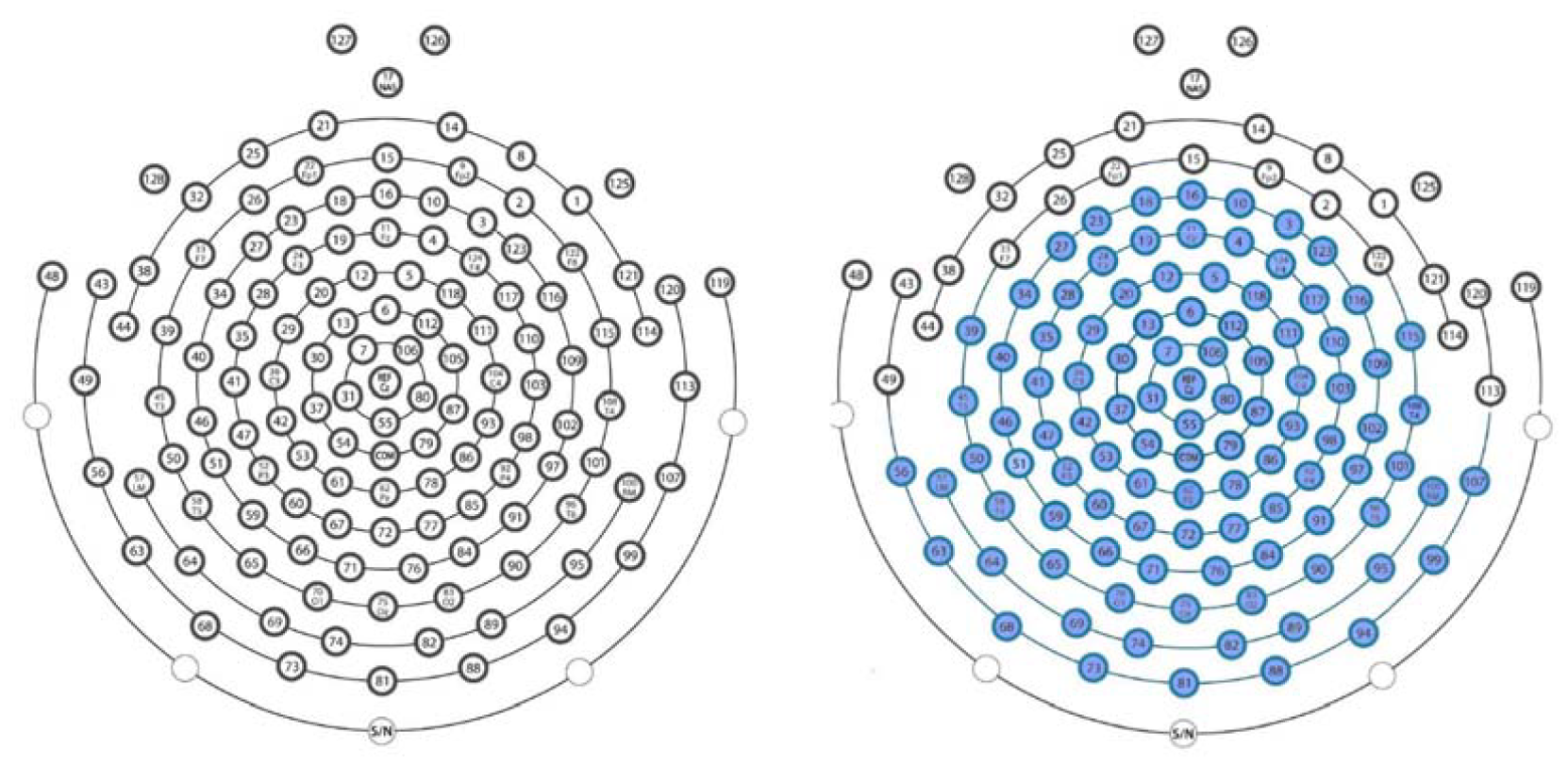
The image on the left depicts the standard, or short, 128 channel infant net, which contains 128 electrodes and pedestals that are all 6.5 mm tall and made with a soft plastic material. The image on the right depicts the modified tall net, where the electrodes shaded in blue are 9.3 mm tall and made of a more rigid plastic material.

**Figure 2.**
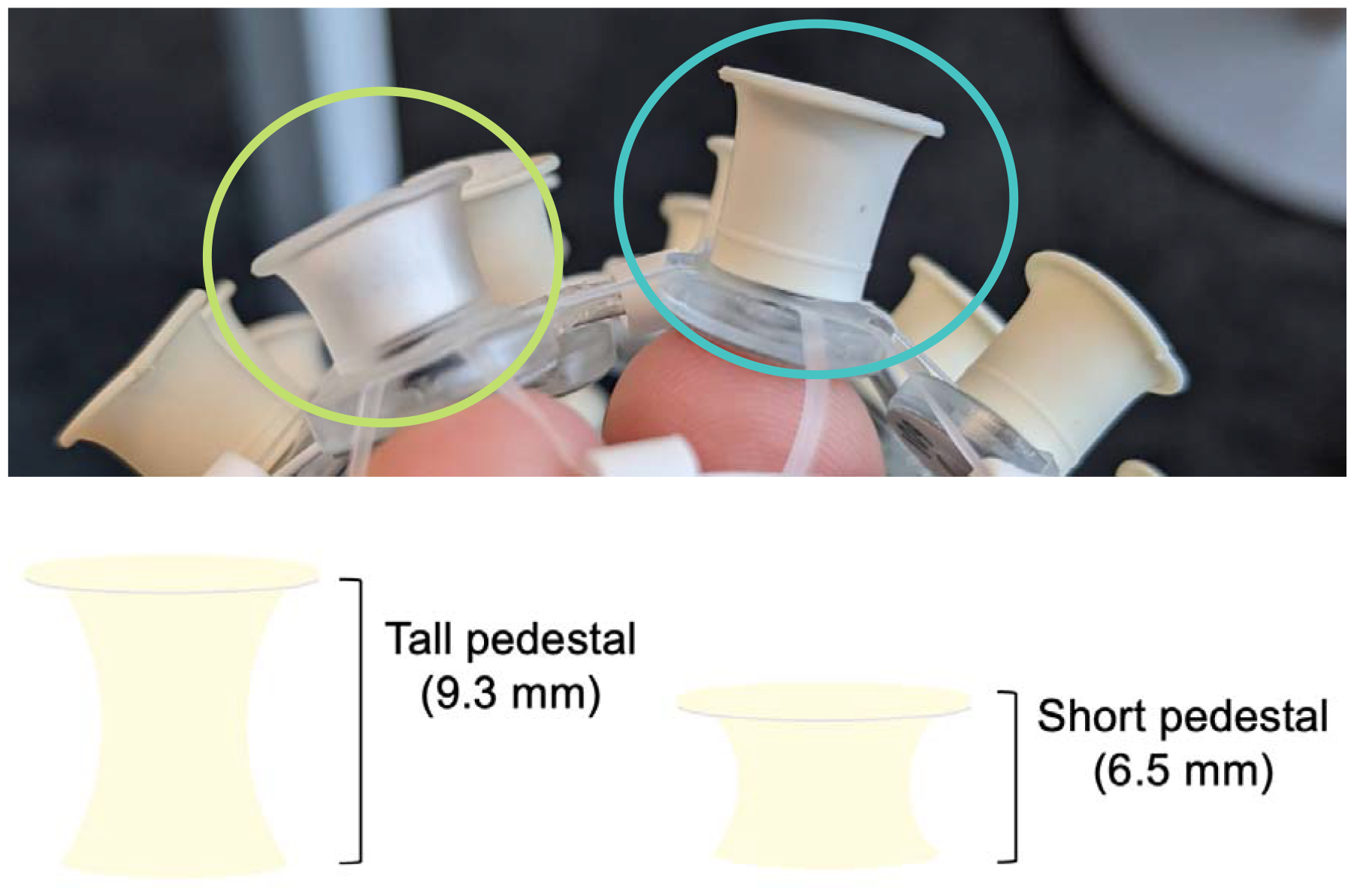
Circled in green (left) is an example of the standard, or short, electrodes of 6.5 mm height from the infant net which uses a clear soft pedestal. Circled in blue (right) is an example of the modified, tall, electrodes of 9.3 mm height and more rigid molded rubber material.

**Figure 3.** Image depicting the modified tall net structure on a child.

In order to best compare the EEG net structures, infants completed two resting state EEG recordings in succession, one recording per net. Prior to net placement, a widely available hair conditioner (SheaMoisture Jamaican Black Castor Oil Strengthen and Restore Leave-In Conditioner) was applied on the infant’s scalp and hair to prevent hair tangling around the pedestals during net placement and removal. This leave-in conditioner contains insulating ingredients so there is minimal risk of electrical bridging, and this solution has not been found to disrupt the recorded EEG signal or the integrity of the net during testing over six years (unpublished data). Conditioning hair in this way allows for the nets to lay closer to the scalp, which will also improve data quality during collection. The use of this product in youth with coarse, curly or tightly coiled hair also speaks to the issue of participant equity in participating comfortably, since this product makes net placement and removal significantly easier by reducing pulling, tugging, and tangling with these hair types.

Resting state EEG data was recorded in a dimly lit room while participants were seated on their caregivers’ lap observing a monitor displaying a brief video. The resting state EEG video was presented via E-Prime software and consisted of a 3-minute long silent video of different colorful and engaging clips. An experimenter was seated beside the infant to help keep them calm and would engage the infant with bubbles or another silent toy as needed during the recording sessions. Data were collected using Net Station 5.4 software and a Magstim EGI Net Amps 400 Series high-input impedance amplifier. Electrode impedances were kept below 100 KΩ when possible in accordance with the amplifier’s impedance capabilities and data was collected at a sampling rate of 1000 Hz with an online reference to channel Cz. Impedance values were run prior to the start of data collection for each net and saved from the Net Station 5.4 recording for analyses.

### 2.3 EEG Data Processing

All EEG files were processed using the Harvard Automated Processing Pipeline for EEG, an automated preprocessing pipeline designed for infant EEG data (HAPPE; Gabard-Durnam et al., 2018; Monachino et al., 2022). Version 4.0 of the HAPPE pipeline was run using MATLAB version 2022b and EEGLAB version 2022.0 (Delorme & Makeig, 2004). Pre-processing parameters are available in Supplemental Table 1. The following electrodes (outer rim of the net structure and eye electrodes removed from the standard net structure for infant comfort) were excluded from further analysis: E125, E126, E127, E128, E48, E119, E43, E49, E56, E63, E68, E73, E81, E88, E94, E99, E017, E113, E120, E44, E38, E32, E25, E21, E14, E8, E1, E121, E114, and E17, which is common practice EEG research with infants (Monachino et al., 2022), leaving 99 remaining electrodes included in analyses. CleanLine (Mullen, 2012) was used to remove 50 Hz electrical line-noise via a multi-taper regression approach, and data were then filtered 1-100 Hz with a finite impulse response (FIR) bandpass filter. Bad channels were detected using HAPPE’s automated algorithm. Data were then subjected to wavelet-thresholding and HAPPE’s MUSCIL (ICA with automated rejection of muscle-contaminated components using EEGlab’s IC-Label algorithm) to correct artifacts. EEG data was segmented into contiguous 2-second windows. Any segments with remaining artifact were removed using amplitude based (+/- 200 uV) and joint probability criteria. Bad channels were then interpolated via spherical spline interpolation and data were re-referenced to the average reference.

### 2.4 Statistical Analysis

Statistical analyses were conducted using SPSS version 28 (IBM Statistics). Primary evaluation outcomes of interest between the two net structures were 1) impedance values obtained prior to each recording both across the online reference electrode (Cz), and all electrodes used in analyses, 2) the percent of good channels in recordings that were retained after pre-processing, and 3) the percent of usable data segments retained after pre-processing. For each of these outcomes of interest, paired samples *t*-tests were performed to compare tall and short net values on each outcome metric within participants. Next, to test whether results for these primary outcomes varied by hair height within the sample, participants were split into two subgroups: 1) 0 cm ≤ 1 cm (*N* = 14) and 2) greater than 1 cm (*N* = 21). The threshold of 1 cm was selected given the sample distribution and the short net sensor height of 6.5 mm, effectively partitioning participants by whether their hair was shorter/close to the standard sensor height, or much taller (almost all more than double) than standard sensor height. Two-way mixed ANOVAs were performed to compare outcome metrics between participant hair height sub-groups.

## 3. Results

### 3.1 Primary Analyses

#### 3.1.1 Online reference electrode impedances

Paired samples *t*-tests examined the impedance values of the online reference electrode (Cz) in short compared to tall net conditions. The analysis revealed a statistically-significant difference in the impedance value of the online reference electrode between short (*M* = 15.52, *SD* = 12.04) and tall conditions (*M* = 10.13, *SD* = 9.02); *t*(34) = 2.69, *p* = .011, *η*^*2*^= .175). Specifically, the tall nets were found to have significantly lower impedances in the online reference electrode than short nets (Figure 4).

**Figure 4.**
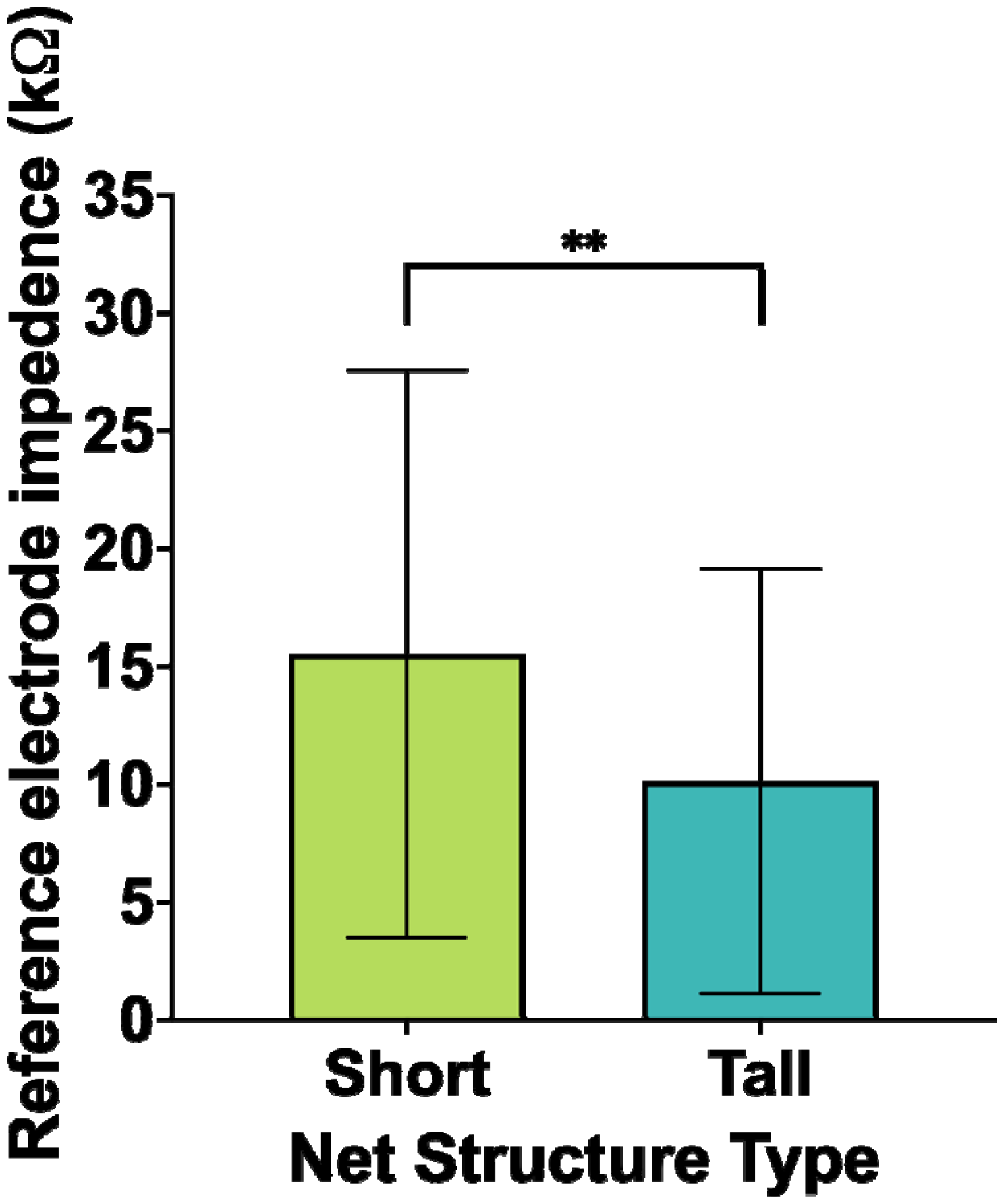
Impedance value of the online reference electrode.

#### 3.1.2 Average electrode impedances

Paired samples *t*-tests examined average impedance values across all electrodes included in pre-processing (i.e., excluding rim electrodes) in short compared to tall net conditions. The analysis revealed a statistically-significant difference in the average impedance value between short (*M* = 17.23, *SD* = 15.04) and tall conditions (*M* = 8.59, *SD* = 7.05); *t*(34) = 3.76, *p* < .001, *η*^*2*^= .294. Specifically, the tall nets had significantly lower average electrode impedances than the short nets for channels retained in pre-processing (Figure 5).

**Figure 5.**
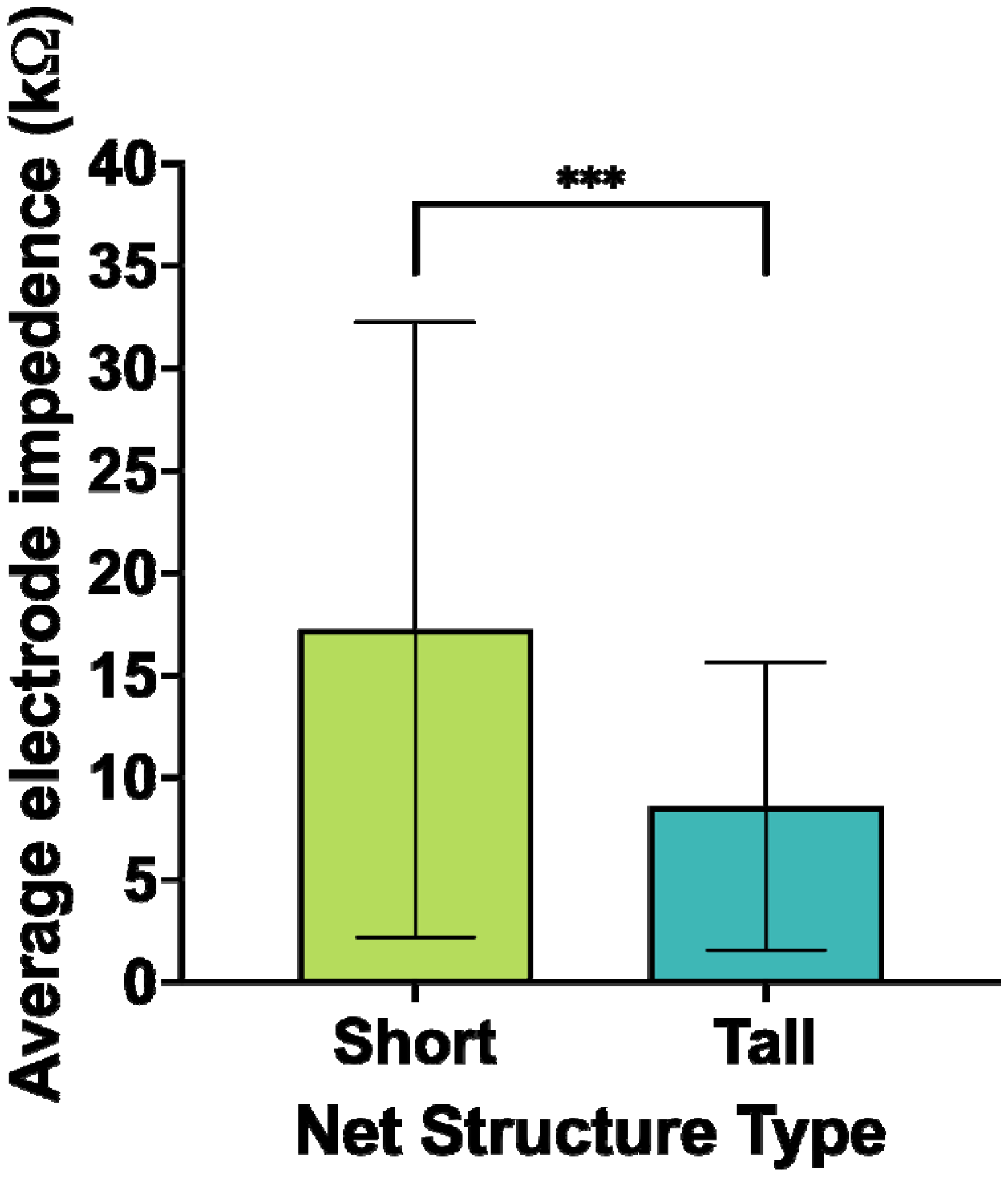
Average impedance value of electrodes used in analyses.

#### 3.1.3 Percent good channels retained

Paired samples *t*-tests examined the percent of good channels retained during recordings in short compared to tall net conditions. The analysis revealed a statistically-significant difference in the percent of good channels kept between short (*M* = 91.98, *SD* = 3.72) and tall conditions (*M* = 95.38, *SD* = 2.71); *t*(34) = -4.28, *p* < .001, *η*^*2*^= .35. Specifically, the tall nets were found to keep a significantly greater percentage of usable channels than the short nets (Figure 6).

**Figure 6.**
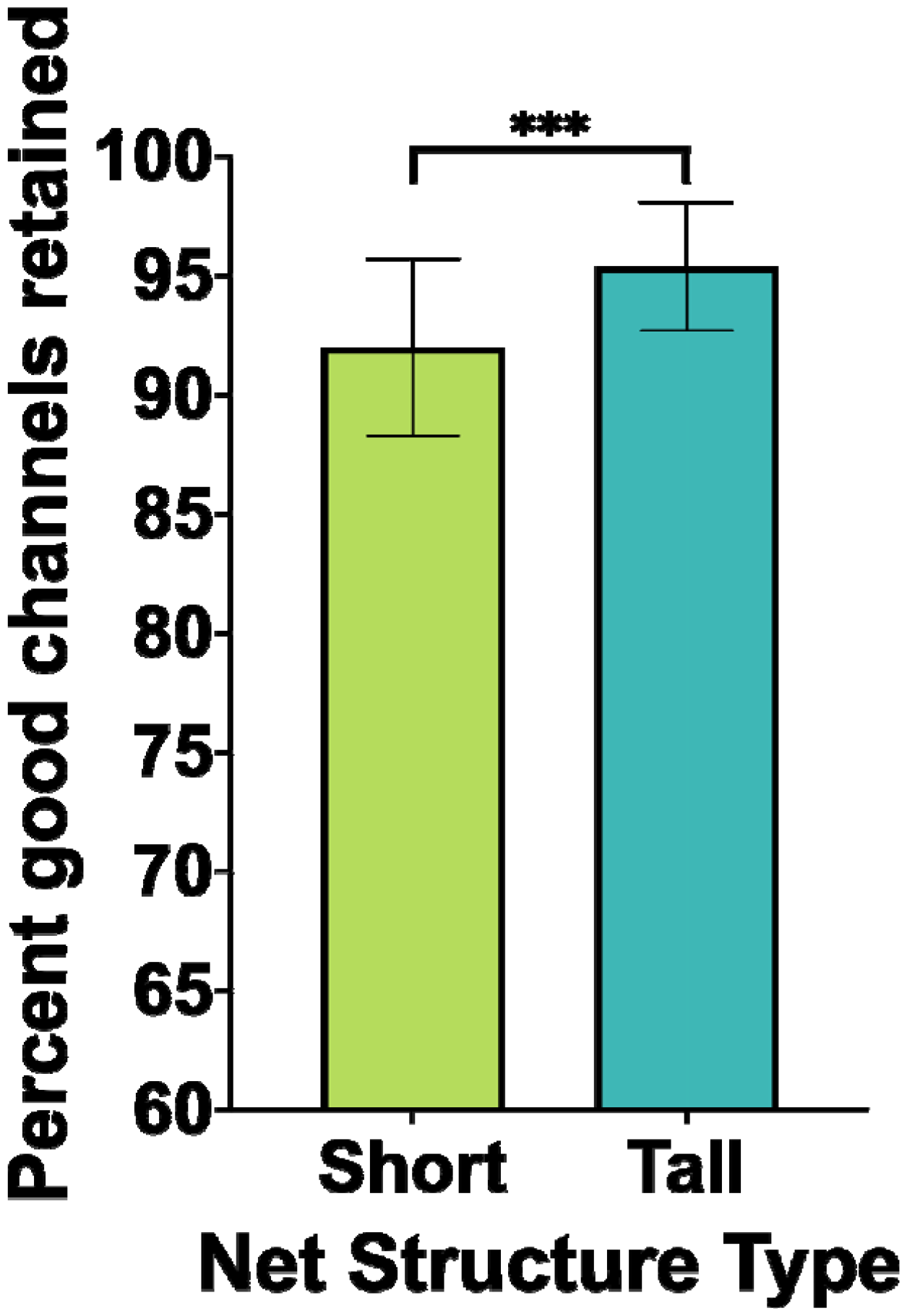
Percent good channels retained post-processing.

#### 3.1.4 Percent usable segments retained

Paired samples *t*-tests examined the percent of usable segments retained during recordings in short compared to tall net conditions. The analysis revealed a statistically-significant difference in the percent of usable segments kept between short (*M* = 68.15, *SD* = 11.19) and tall conditions (*M* = 73.55, *SD* = 6.22); *t*(34) = -2.77, *p* = .009, *η*^*2*^= .184. Specifically, the tall nets were found to keep a significantly greater percentage of usable segments for analysis than the short nets (Figure 7).

**Figure 7.**
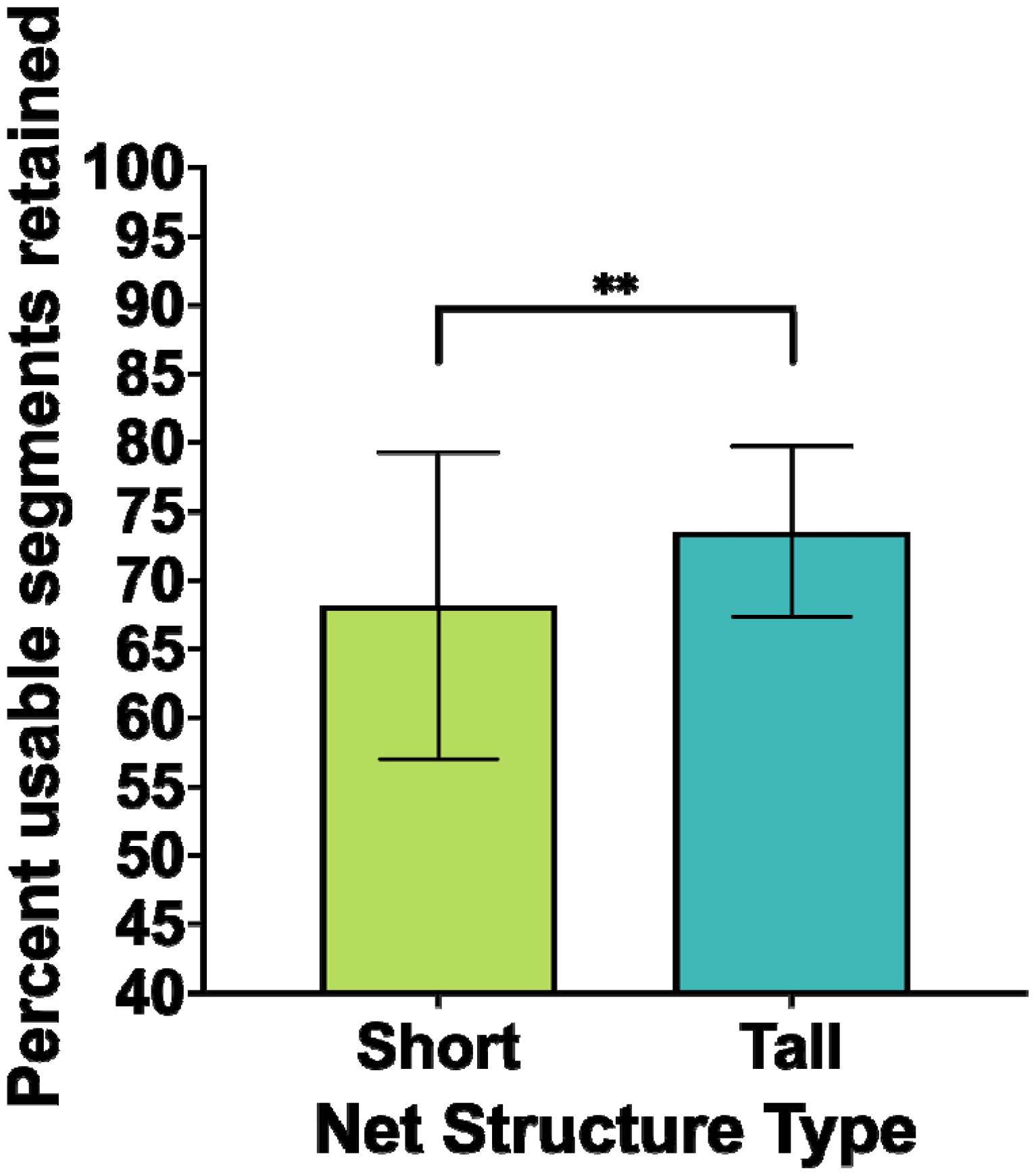
Percent usable segments retained post-processing.

### 3.2 Analyses by Hair Height

#### 3.2.1 Online reference electrode impedance

Two-way mixed ANOVAs examined the effect of hair height on net performance. We do not find a statistically-significant interaction between net pedestal height and hair height group in predicting the impedance value of the online reference electrode, *F*(33) = 0.41, *p* = .530,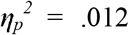 . However, exploratory comparisons revealed that while there are no significant mean differences observed between net pedestal height on electrode impedance for those with hair shorter than 1 cm, the impedance values for the online reference were significantly higher for the short net (*M* = 17.95, *SE* = 2.58) than the tall net (*M* = 11.51, *SE* = 1.96; *p* = .019, 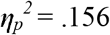) when participants had hair greater than 1 cm in length. (Figure 8).

**Figure 8.**
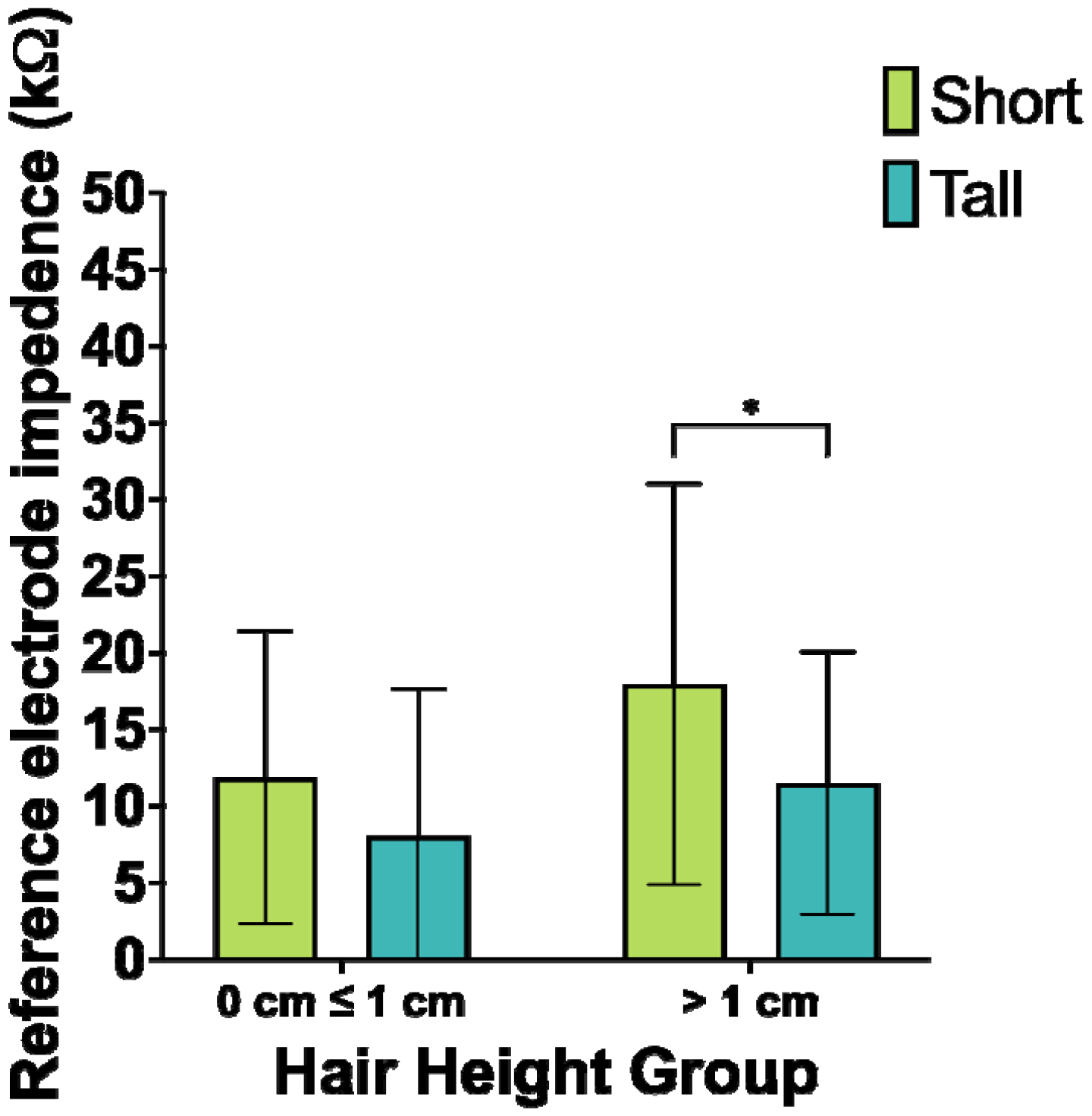
Impedance value of the online reference electrode by hair height.

#### 3.2.2 Average electrode impedance

wo-way mixed ANOVAs examined the effect of hair height on net performance. We did not find a statistically-significant interaction between net pedestal height and hair height group in predicting percent average impedance values, *F*(33) = .09, *p* = .756,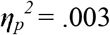.

#### 3.2.3 Percent good channels retained

Two-way mixed ANOVAs examined the effect of hair height on net performance. We did not find a statistically-significant interaction between net pedestal height and hair height group in predicting percent of good channel remaining after processing, *F*(33) = 0.25, *p* = .622, 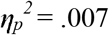.

#### 3.2.4 Percent usable segments retained

Two-way mixed ANOVAs examined the effect of hair height on net performance. We did not find a statistically-significant interaction between net pedestal height and hair height group in predicting percent of usable segments remaining after processing, *F*(33) = 1.09, *p* = .304, 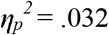.

## 4. Discussion

The current study compared EEG data quality and retention between two high-density net designs from the same company, Magstim EGI to evaluate a potentially more inclusive net design for use with infants and toddlers across hair types and hair height. We found that the modified net with taller, 9.3 mm pedestals on a portion of the scalp performed better overall than the short nets in keeping a higher percentage of good channels and usable segments, as well as lowering impedances in infants 6–11 months old with curly and/or tightly coiled hair types. Further, the modified net with tall pedestals specifically performed better than the short net at lowering impedances in the online reference electrode for infants with hair greater than 1 cm in length in this sample. Overall, using the modified tall pedestal net, as compared to the short net, resulted in higher levels of data retention, better signal quality, and better spatial distribution of usable data in infancy for those with curly, tightly coiled, or tall hair.

### 4.1 Limitations

The current study has several limitations to consider. First, the sample size for analysis was small, however the within-person design is a strength in addressing the aims of this study. Testing order of the nets is also a limitation, as for all the infants the short net was placed first and then removed and the tall net was placed last, although the temporal delay between placements was only approximately 5 minutes between recordings. Considering more limited participant tolerance and attention in infancy, this may have had an impact on the overall findings, as infants may have been fussier for the second round of recording compared to the first recording. However, this would likely lead to underestimates of the effect of the net design change as fussier infants would not contribute higher data quality. Conversely, the tall net impedance measurements may have benefited from pre-wet scalps, though this is unlikely to have produced the full results set observed, including the hair-height specific results. Additionally, measuring hair height from one scalp location in infancy is an incomplete measure of hair height as hair emerges and grows inconsistently across the scalp early in life. Thus, while the hair height measures used in this study do not necessarily index hair for the entire scalp, they did provide a consistent reference for variation at one scalp location across the infants sampled here. Moreover, this simple height measurement doesn’t account for hair density and how that property could potentially affect EEG signal quality, which remains an important unanswered question in this research space. Lastly, a leave-in conditioner was applied prior to placing the nets for all the infants. This solution has not been found to disrupt the recorded EEG signal or the integrity of the net during testing over six years (unpublished data), but does reduce hair height, so this practice might impact findings in the current study by underestimating differences between nets that would be clearer without the use of conditioner. The conditioner was included for all babies here given our previous observations that it improves participant comfort with the hair types in this sample (see recommendations below).

### 4.2 Recommendations

Based on our testing, we provide recommendations for using the modified tall-pedestal net design and improving the inclusivity of EEG research with infant and toddler participants with curly or tightly-coiled hair. First, regardless of net type used, we recommend using a widely-available conditioner (e.g., SheaMoisture Jamaican Black Castor Oil Strengthen and Restore Leave-In Conditioner; Cantu Shea Butter Leave In Conditioning Repair Cream) on participant’s scalp and hair prior to net placement and data collection for saline-based high-density EEG recordings. The use of this product helps to reduce tangling during net placement and net removal and provides additional moisture that can protect hair from drying out from the saline solution. This detangling and moisturizing aspect of adding conditioner can also make it easier for electrodes to sit flush against the scalp for recording. Second, when placing the tall-pedestal caps, the tall pedestal electrodes are more likely to be tilted initially instead of sitting flush on the scalp. When possible, an extra pair of hands from a researcher on the study or from the participant’s caregiver to help align electrodes perpendicular to the scalp will be useful in reducing the time it takes to place the EEG net. Finally, the modified tall pedestals impacted tension and sizing in the initial net fit, so we sized down the tall pedestal caps by 1 cm (e.g., 44 - 47 cm range becomes 43 - 46 cm) in order to ensure that the cap was secure and the electrodes sat flush against the scalp for better quality recordings. This is now reflected in Magstim EGI’s accurate sizing of the tall pedestal net as a commercially available product.

### 4.3 Conclusion

Here we found evidence that a modified net design with taller EEG pedestals produced more and higher-quality EEG data than the standard net design in infants with curly, tightly-coiled, or tall hair height. This modified net design is now commercially available and we recommend its use in infant studies moving forward. These findings are specific to the Magstim EGI infant sized nets, and future research will benefit from exploring other net configurations from Magstim EGI for older ages (where configurations of pedestal length and material differ across the scalp by age). In sum, in conjunction with other best practices, the modified tall-pedestal infant net design is a promising technological modification to improve the data quality, data retention, and inclusivity of EEG research with infant and toddler participants with curly or tightly coiled hair.

## Acknowledgements

We would like to thank the families in the Khula study for their participation that made this work possible. We would also like to thank the recruitment team for the Khula study, especially Ringie Gulwa and Pamela Madikane. Thank you also to Jordan Bucenec, Maille Hagen, Finn Janak, Elena Lam, Oluwafunso Olaniyan, and Ana Sobrino for their assistance with data organization. Thank you to Saniya Burman and Emma Margolis for their assistance with figure design. Thank you to members of the PINE Lab at Northeastern and the Khula study staff at the University of Cape Town for relevant discussion. This research is supported by funding from the Wellcome Leap 1kD Program to KAD and LJGD.

---

**Supplemental Table 1.**
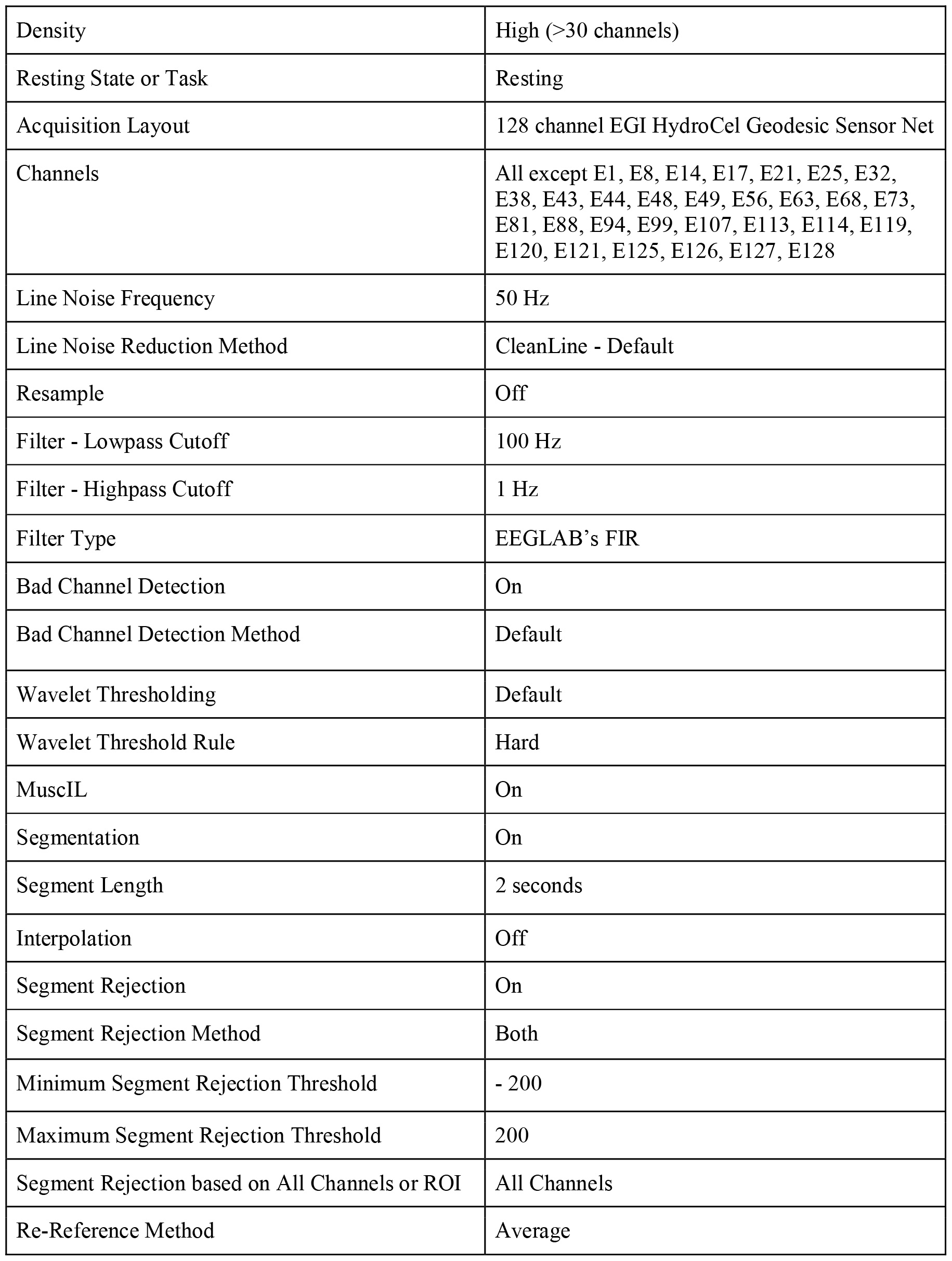
Parameters used in HAPPE processing.

## Notes

### Competing Interest Statement

The authors have declared no competing interest.

